## Supplementary Information for "CUT&Tag for efficient epigenomic profiling of small samples and single cells"

**Data processing for all Figures in Kaya-Okur *et al* (2019).**

**Peak calling:**
In all cases except as noted MACS2 was used to call peaks on bedfiles for CUT&RUN and CUT&Tag,datasets with no input files with the following flags:
macs2 callpeak -t input_BED -f BEDPE -p 1e-5 --keep-dup all -n output_prefix

**Heatmapping:**
In all cases the midpoints of peak regions were used as the center in +/- 1kb regions with 50 bp bins were generated, ordered by mean signal and plotted using deepTools. The display range was adjusted to the maximum and minimum signals for each heatmap, and the number of regions is reported in the figure.

**New data:**
All sequencing data is deposited in GEO under the super-entry GSE124690, including the bulk GSE124557 and the single-cell entries GSE124680 and GSE124683.

**Software packages used:**
MACS2 version 2.1.1, deepTools version 3.0.2, seaborn version 0.8.1, and FIMO version 5.0.4.

**For Figure 2a:**
UCSC Genome Browser tracks for datasets (GSM788088, GSM3391661, GSM3536515, GSM3536516, GSM3536517, GSM3536520, GSM3560264, GSE99173) from K562 cells were plotted with autoscaling, except for the IgG control, which was set to a track height of 20.

**For Figure 2b:**
UCSC Genome Browser tracks for datasets (GSM733748, GSM3536498, GSM3536499, GSM3536500, GSM3536502, GSM3560256) from H1 cells were plotted with autoscaling, except for the IgG control, which was set to a track height of 20.

**For Figure 2c:**
K4me1 bedfiles for CUT&RUN (GSM3391662) and for CUT&Tag (GSM3536516) were used for heatmapping. For H3K4me1 ChIP-seq, we used the list of peaks associated with the GEO entry GSM733692.

**For Figure 2d:**
Bedfiles for RNAPII CUT&Tag (GSM3391654) were used for heatmapping. Promoters were ordered by gene expression determined by RNA-seq (GSM207236). Promoters within 1 kb of each other were excluded. This promoter list is provided in supplementary table 1.

**For Figure 2e:**
Peaks for RNAPII-S5p CUT&Tag were called using MACS2 on the HsSc_PolS5_F_0802, HsSc_PolS5_T_0802 dataset. PRO-seq data (SRA GSM1480327) for K562 cells were clustered using a custom script described in Ref. 10. Heat maps were displayed using Java Treeview (http://jtreeview.sourceforge.net/).

**For Figure 2f:**
Bedfiles for ATAC-seq (GSM2695560) and for H3K4me2 CUT&RUN (GSM3391663) were used for peak calling. For ATAC calls, we performed a tail truncation removing the 2.5% most

significant and least significant peaks to limit sites enriched for PCR duplicates or with very low read counts. Heat maps were then plotted.

**For Figure 3a:**
We divided the genome into 500 bp bins and reads in each bin from H3K4me1 for ChIP-seq (GSM733692), CUT&RUN (GSM3391662), and CUT&Tag (GSM3536516) experiments along with CUT&Tag for IgG (GSM3560264) were summed. Pearson correlations were generated by comparing between all pairs of the log transformed read count array. These Pearson correlations were then used for hierarchical clustering and the figure was generated using the seaborn package on Python.

**For Figure 3b:**
We subsampled each of the datasets to 20M, 10M, 5M, 2.5M, 1M, and 500K mapped reads, and then called peaks on each set. For ChIP-seq and ATAC-seq datasets, we changed the flag -f to BED instead of BEDPE because these datasets were single-end reads. The sets of peaks for each subsampled population of reads are provided in Supplementary table 1. We then binned the reads in a +/- interval around peaks and calculated the percent of reads in peaks of the total read count.

**For Figure 4a:**
IGV Genome Browser track for NPAT CUT&Tag data (GSM3536519) for chromosome 6 was plotted with autoscaling.

**For Figure 4b:**
To determine read counts at histone gene promoters, we binned read counts within a +/- 100 bp region of each histone gene in a NPAT CUT&Tag experiment (GSM3536519). To determine read counts at hypersensitive sites, we repeated binning and counting for all ATAC-seq peaks that do not overlap with a histone gene promoter. To determine background, we selected 10,000 positions in the genome at random and counted reads at those sites. These read counts are plotted in the histogram.

**For Figure 4c:**
We curated a list of replication-dependent histone genes and counted NPAT CUT&Tag reads mapping around +/- 100 bp of the summit of their promoters. We generated a list of “ATAC” sites as peak calls from ATAC-seq data, and then removed histone gene promoters from that list. We then counted reads that mapped +/- 100 bp of the summit of these “ATAC-only” sites, and generated a histogram to display the results.

**For Figure 5a:**
We obtained the MA0139.1 CTCF motif for humans from the JASPAR database. We used FIMO (http://meme-suite.org/doc/fimo.html) with default parameters to search the hg19 genome for matches to the CTCF motif, and ordered sites by their p-value. Read counts from CTCF ChIP-seq (**GSM733719**), CUT&RUN (GSM3391659), and CUT&Tag (GSM3560258) experiments on this list of sites were heatmapped.

**For Figure 5b:**
We generated a list of CTCF-bound sites by calling peaks on CTCF CUT&RUN data (GSM3391659), and counted CTCF CUT&Tag reads mapping around +/- 100 bp of the summit of those sites. We generated a list of “ATAC” sites as peak calls from ATAC-seq data, and then removed CTCF-bound sites from that list. We then counted reads that mapped +/- 100 bp of the summit of these “ATAC-only” sites, and generated a histogram to display the results.

**For Figure 5c:**
We used the list of CTCF motifs from Figure 5a, and mapped read ends centered on these motifs for each salt condition (GSM3560257, GSM3560258, and GSM3560259). We then smoothed mapped read ends along a 11bp window using a Savitzky-Golay filter (SciPy) with a polynomial order of zero.

**For Figure 5d:**
UCSC Genome Browser tracks for CTCF CUT&Tag (CTCF_5m_1), ATAC (ATAC_5m_1), H3K4me2 ChIP-seq (ENCODE_5m_1), H3K4me2 CUT&Tag (H3K4me2_5m_1), H3K4me3 CUT&Tag (H3K4me2_5m_2), H3K4me1 CUT&Tag (H3K4me1_5m_1), H2A.Z CUT&Tag (H2AZ_5m) were plotted with autoscaling.

**For Figure 6:**
IGV Genome Browser tracks for 908 individual single cells profiled by H3K27me3 scCUT&Tag (GSM124680) and for 807 individual single cells profiled by H3K4me2 scCUT&Tag (GSM124683) were displayed showing individual fragments.

**For Supplementary Figure 1A:**
Read counts mapping to the hg19 and *E. coli* genomes from a serial dilution of starting cells (HK_Hs_K5II_K27me?_*_0912) is plotted.

**For Supplementary Figure 1B:**
IGV Genome Browser tracks for H3K27me3 CUT&Tag using different starting cell numbers (HK_Hs_K5II_K27me5_6k_0912.bed, HK_Hs_K5II_K27me7_600_0912.bed, and HK_Hs_K5II_K27me9_60_0912.bed) was plotted with autoscaling.

**For Supplementary Figure 2A:**
UCSC Genome Browser tracks for (GSM3536521, GSM3536522, GSM3536523, GSM3536518, GSM3536514) were plotted with autoscaling.

**For Supplementary Figure 2B:**
UCSC Genome Browser tracks for (GSM3536501, GSM3536497, GSM3536506) were plotted with autoscaling.

**For Supplementary Figure 3A:**
UCSC Genome Browser tracks for (GSM3536521, GSM3536522, GSM3536523, GSM3536518) were plotted with autoscaling.

**For Supplementary Figure 3B:**
Bedfiles for RNAPII CUT&Tag (GSM3536521, GSM3536522, GSM3536523) were used for heatmapping. Promoters were ordered by gene expression determined by RNA-seq (GSM207236). Promoters within 1 kb of each other were excluded. This promoter list is provided in Supplementary Table 1.

**For Supplementary Figure 3C:**
We ordered gene promoters by RNA-seq (Supplementary Table 1) and plotted read counts on (+) and (–) strands from RNAPII S2p (GSM3536521), RNAPII S5p (GSM3536522), and RNAPII S7p (GSM3536523) CUT&TAG experiments. Strands were recolored in Photoshop and levels were adjusted to maximize display intensity.

**For Supplementary Figure 4:**
Unique mapped read counts for 908 individual single cells profiled by H3K27me3 scCUT&Tag (GSM124680) and for 807 individual single cells profiled by H3K4me2 scCUT&Tag (GSM124683) were graphed.

**For Supplementary Figure 5A:**
Library profiles after CUT&Tag experiments were measured using an Agilent Tapestation 4200 with D1000 High Sensitivity Screen tapes.

**For Supplementary Figure 5B:**
IGV Genome Browser track for CUT&Tag with the following datasets were plotted with fixed scaling: SH_Hs_NPA1_20190226**,** SH_Hs_NPA2_20190226**,** SH_Hs_NPA4_20190226, SH_Hs_NPA8_20190226, SH_Hs_NPB1_20190226, SH_Hs_NPB2_20190226, SH_Hs_NPB4_20190226, SH_Hs_NPB8_20190226, SH_Hs_me3A1_20190226, SH_Hs_me3A2_20190226, SH_Hs_me3A4_20190226,SH_Hs_me3A8_20190226, SH_Hs_me3B1_20190226, SH_Hs_me3B2_20190226, SH_Hs_me3B4_20190226, and SH_Hs_me3B8_20190226.

**Software References**

deepTools:
Ramírez, F. *et al*. deepTools2: a next generation web server for deep-sequencing data analysis. *Nucleic Acids Research* **44**, W160-W165 (2016).

MACS2:
"Zhang, Y. *et al*. Model-based Analysis of ChIP-Seq (MACS). *Genome Biology* **9**, R137 (2008).

FIMO:
Grant, C.E. *et al*. FIMO: scanning for occurrences of a given motif. *Bioinformatics* ***27***, 1017-1018 (2011).

JASPAR:
Khan, A. *et al*. JASPAR 2018: update of the open-access database of transcription factor binding profiles and its web framework. *Nucleic Acids Research* **46**, D260–D266 (2018).

RNA-seq data:
ENCODE Project Consortium. An integrated encyclopedia of DNA elements in the human genome. *Nature* **489**, 57-74 (2012).
